# Prime Editing Models the MTARC1 A165T Variant in Human Liver Organoids, Demonstrating Reduced Steatosis, Inflammation, and Fibrosis

**DOI:** 10.64898/2025.12.23.696200

**Authors:** Amel Ben Saad, Arden D Weilheimer, Stefan D Gentile, Nahid Arghiani, Benjamin J Toles, Sudipta Tripathi, Seher Mohsin Sayed, Anil Chandraker, Alan C Mullen

## Abstract

Metabolic dysfunction-associated steatotic liver disease (MASLD) is the most prevalent cause of chronic liver disease. MASLD is a progressive and multifactorial disease marked initially by hepatic steatosis, which can progress to steatohepatitis, fibrosis, cirrhosis, and liver cancer. Genetic factors influence the development, progression, and complications in MASLD, and genome-wide association studies (GWAS) have identified single nucleotide polymorphisms (SNPs) associated with altered risk. Mitochondrial amidoxime reducing component 1 (*MTARC1*) *rs2642438* (p.A165T) variant has been identified as protective, but the role of *MTARC1* and the impact of this variant in hepatocytes remains poorly understood. Here, we applied prime editing to create the *rs2642438* variant in human pluripotent stem cells (hPSCs) before differentiation into human liver organoids (HLOs) to investigate the effect of the variant under conditions of steatotic and fibrotic injury. Compared to HLOs formed from hPSCs containing the *MTARC1* reference sequence, HLOs with the *rs2642438* variant show lower levels of MTARC1 protein and triglycerides and are protected from steatotic and fibrotic injury, as predicted by the phenotype observed in patients carrying the variant. The observed decrease in triglyceride level with the variant appears to be driven more by suppression of de novo lipogenesis rather than stimulation in ß-oxidation in the HLO model. While resmetirom, the thyroid hormone receptor-beta (THRB) agonist approved to treat patients with metabolic dysfunction-associated steatohepatitis (MASH) was effective in reducing triglyceride levels in the setting of steatotic injury in HLOs with the reference sequence, HLOs containing the variant did not show further reduction in triglyceride levels with exposure to resmetirom, despite increased expression of *THRB*. Together, this study establishes an approach to model disease-related SNPs in HLOs and provides further insights into the activity of the *MTARC1* variant, and suggests that profiling SNPs may be a path to identify patients more likely to respond to therapies for MASLD.

## INTRODUCTION

Chronic liver disease (CLD) remains a major global health concern, causing more than two million deaths annually (1) and affecting approximately 4.5 million adults in the U.S. (2). The most common cause of CLD is metabolic dysfunction-associated steatotic liver disease (MASLD). MASLD affects more than 30% of the adult population, and its prevalence is predicted to increase in the coming years (3). Furthermore, a concerning rise in MASLD prevalence has been observed among young children and adolescents, with current estimates indicating nearly one in ten children in the U.S. are affected (4). MASLD is a progressive disease, starting with steatosis, which can progress to metabolic dysfunction-associated steatohepatitis (MASH), fibrosis, cirrhosis, and liver cancer (5, 6). Therapeutic options for patients with MASLD are limited. Lifestyle improvement including weight loss, Mediterranean diet, limiting alcohol consumption, and physical exercise are first-line therapies (7). Recently, resmetirom, a thyroid hormone receptor-B (THRB) agonist, was approved for adults with MASH and moderate fibrosis (8), and the GLP-1 receptor agonist semiglutide has now been shown to reduce fibrosis in MASH (9). While these treatments are significant advances, only 25% of patients receiving resmetirom show improved fibrosis (10), and about a third of patients show improvement in the combined endpoints of steatohepatitis or fibrosis with semiglutide (9). A better understanding of the pathogenesis and progression of MASLD is still needed to identify more effective treatments.

MASLD is a complex and multifactorial disease. Different factors can regulate its development, progression, and complication, including genetic factors (11). Genome-wide association studies (GWAS) have identified susceptibility loci or single nucleotide polymorphisms (SNPs) associated with an altered risk in developing MASLD, including SNPs in patatin-like phospholipase domain-containing 3 (*PNPLA3*), transmembrane 6 superfamily member 2 (*TM6SF2*), glucokinase regulatory protein (*GCKR*), membrane-bound O-acyltransferase domain-containing 7 (*MBOAT7*), hydroxysteroid 17 beta dehydrogenase 13 (*HSD17B13*) and mitochondrial amidoxime reducing component 1 (*MTARC1*) (12). *PNPLA3 rs738409* (p.I148M) is the most common and replicated variant among different populations, and is strongly associated with an increased hepatic triglyceride level, liver fibrosis, and cirrhosis (13). Similarly, *TM6SF2 rs58542926*, *GCKR rs780094,* and *MBOAT7 rs641738* variants are associated with a higher risk for chronic liver disease (14–16). In contrast, *HSD17B13 rs72613567* and *MTARC1 rs2642438* are reported as protective against MASLD and its progression (17–19).

*MTARC1 rs2642438 G>A* (p.A165T) was initially reported by Emdin et al, and was linked to decreased markers of liver injury (18). Patients carrying this variant have lower levels of circulating alanine and aspartate transaminases (ALT and AST) and serum cholesterol, decreased hepatic steatosis and liver fibrosis, lower lobular inflammation, reduced severity and risk of MASH, autoimmune hepatitis, alcoholic liver disease, and all causes of cirrhosis, as well as a lower rate of liver-related mortality (18, 20–23). The *MTARC1* gene (known also as *MARC1* and *MOSC1*) encodes a molybdenum enzyme located in the outer mitochondrial membrane. It reduces a variety of N-oxygenated compounds and plays a role in drug metabolism and cellular detoxification (24, 25). The protective effect of the *MTARC1* p.A165T variant is linked to reduced MTARC1 protein levels in the liver (26, 27). Studies have explored the potential of *MTARC1* depletion to phenocopy the protective effects of the variant, and N-acetylgalactosamine conjugated siRNAs (GalNac-siRNA) targeting *MTARC1* in primary human hepatocytes and diet-induced MASH mouse models attenuate hepatic steatosis and inhibit disease progression (28–30). These findings support the therapeutic potential of targeting *MTARC1* to treat MASLD, and clinical trials have been initiated to investigate *MTARC1* depletion with GalNac-siRNAs (*24*).

While there is strong evidence supporting the hepatoprotective effect of *MTARC1* p.A165T variant and its potential as a therapeutic candidate, the molecular mechanisms by which MTARC1 variants influence the trajectory of disease remain poorly understood. Thus, in vitro human models are needed to explore and understand the implications of the *MTARC1 rs2642438* p.A165T variant in the liver.

In this study, we created a model of the *MTARC1 rs2642438* p.A165T variant using liver organoids (HLOs) derived from human pluripotent stem cells (hPSCs). We edited hPSCs with prime editing before differentiating hPSCs with the *MTARC1* reference or variant sequence into HLOs, a multicellular system, to study their phenotype and response to liver injury in conditions that mimic MASH and fibrotic injury.

## RESULTS

### Generation of *MTARC1* -WT and -A165T hPSCs and differentiation into HLOs

The *MTARC1* variation *rs2642438 G>A* (p.A165T) is a single nucleotide change of guanine (G) to adenine (A), substituting the amino acid alanine (A) for threonine (T) at amino acid position 165. hPSCs cells used in this study were initially heterozygous (A/G) (**Fig. 1A**). We used prime editing to create hPSCs with the homozygous *MTARC1* reference sequence (*MTARC1*-WT, G/G) and hPSCs with the homozygous *MTARC1* A165T variant (*MTARC1*-A165T, A/A). For each *MTARC1* genotype (G/G and A/A), we generated four independent single-cell derived clones with the desired homozygous modification. Successful editing was confirmed by Sanger sequencing (**Fig. 1B**).

**Fig. 1.**
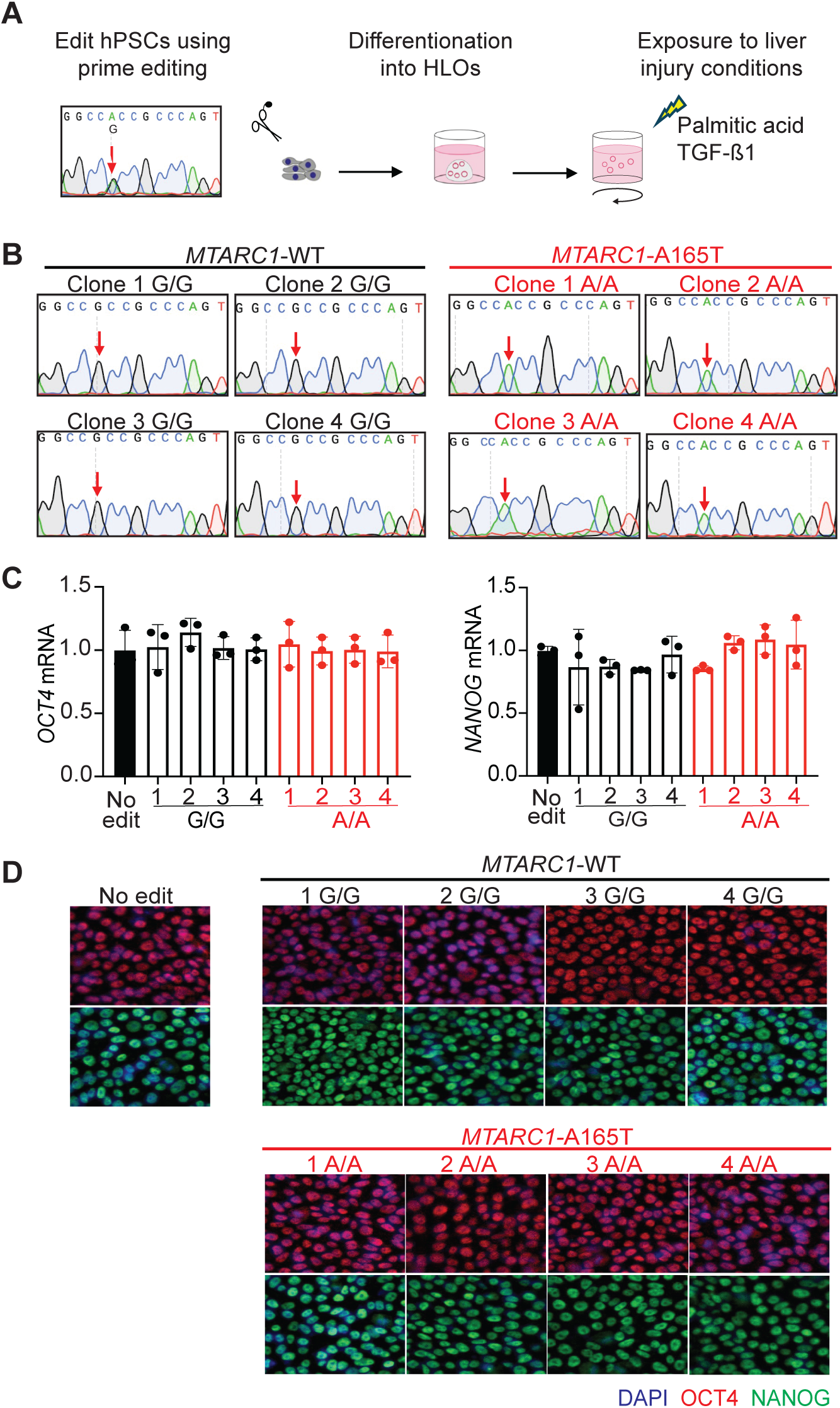
**Generation and characterization of hPSCs with wildtype *MTARC1* and the *MTARC1*-A165T variant**. (A) Study overview and hPSC genotype. hPSCs (A/G) were edited using prime editing to generate *MTARC1*-WT (G/G) and *MTARC1*-A165T (A/A) clones and then differentiated into HLOs. HLOs were extracted from matrigel after maturation (21 days) and maintained in an orbital shaker. HLOs were treated with PA to model MASH and TGF-ß1 to induce fibrosis. (B) Sanger sequencing data for *MTARC1* gene in 8 hPSCs clones after prime editing to confirm the modification. We generated 4 clones for each genotype, G/G genotype (*MTARC1* -WT sequence) (left panel) and A/A genotype (*MTARC1*-A165T variant) (right panel). The red arrows indicate the position of each edit. (C) *OCT4/ POU5F1* and *NANOG* expression levels in hPSCs clones were quantified by RT-qPCR. (D) Immunostaining for OCT4/POU5F1 (red) and NANOG (green) was performed for each hPSC clone. Nuclei are visualized with DAPI (blue). Scale bar, 100 um.

After gene editing, we assessed pluripotency. We found similar expression of the pluripotency markers *NANOG* and *OCT4/POU5F1* in all edited hPSCs, at levels comparable to non-edited hPSCs, as quantified by RT-qPCR (**Fig. 1C**) and immunostaining (**Fig. 1D**). Additionally, hPSC morphology was preserved in the edited hPSC clones; they exhibited typical embryonic stem cell colony morphology (**Fig. 2A**), confirming further that prime editing did not interfere with pluripotency features. We then differentiated all *MTARC1*-WT (G/G) and *MTARC1*-A165T (A/A) hPSC clones to HLOs (31). Regardless of their genotype, all hPSCs clones were successfully differentiated into HLOs when evaluated on day 21 **(Fig. 2A)**.

**Fig. 2.**
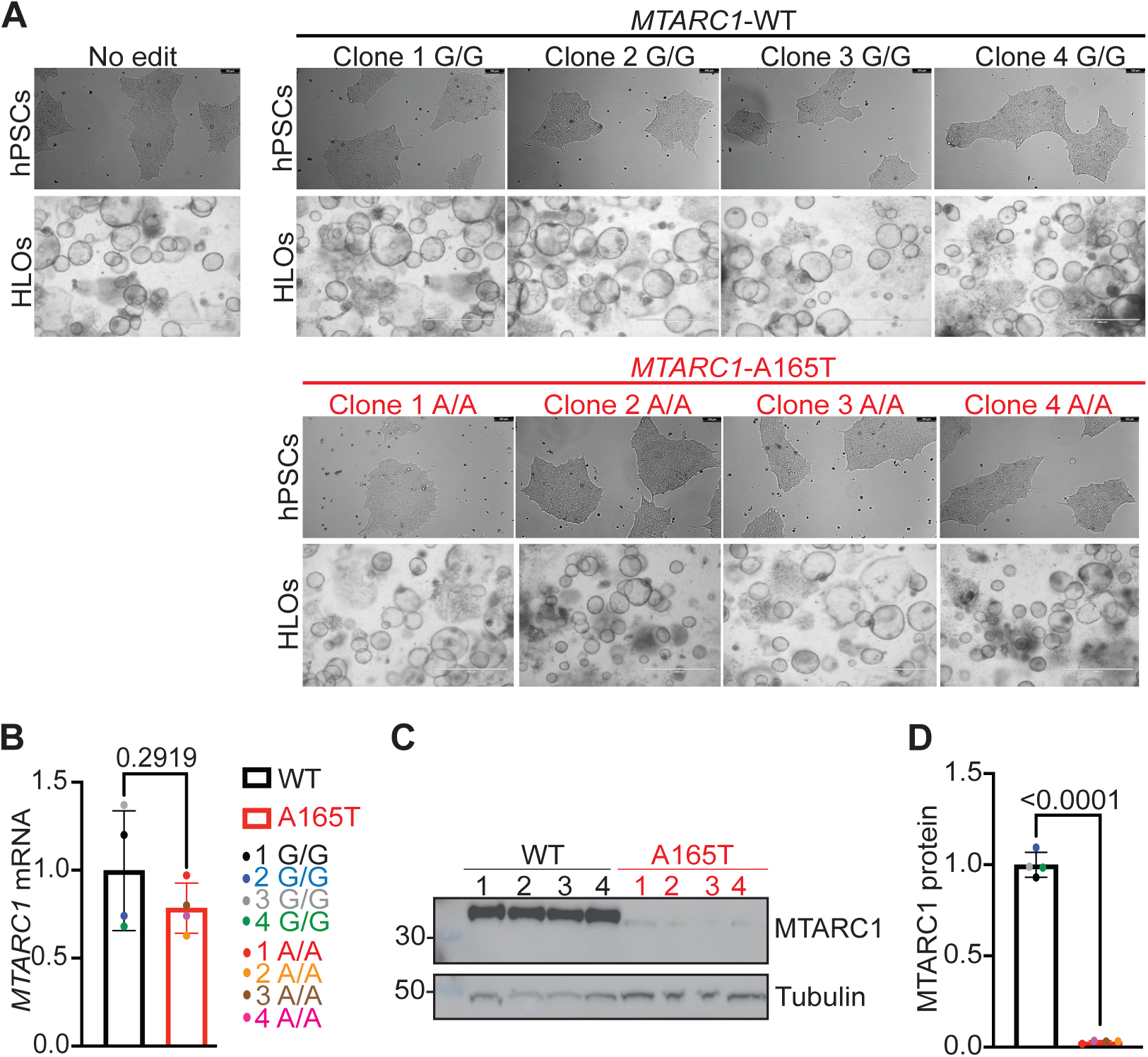
Differentiation of *MTARC1*-WT and -A165T hPSCs into HLOs. (A) Bright field images of hPSC clones (top panel) and HLOs in matrigel on day 21 of differentiation (bottom panel). Scale bar, 100 um. (B) Quantification of *MTARC1* mRNA levels in HLOs carrying *MTARC1*-WT and - A165T by RT-qPCR. Data are representative of 3 independent experiments, and in each experiment, every clone was differentiated in triplicate. P values were calculated using an unpaired two-tailed Student’s t-test. (C) MTARC1 protein levels were quantified by Western blot (top), and tubulin was quantified as a loading control (bottom). (D) MTARC1 band intensities were quantified and normalized to the tubulin signal using Image J software. Data are representative of 3 independent experiments. P value was calculated using an unpaired two-tailed Student’s t-test.

### *MTARC1* A165T variant induces protein loss and reduces lipid accumulation

Following the differentiation of *MTARC1*-WT and -A165T hPSC clones to HLOs, we determined the impact of their genotype at *MTARC1* transcript and protein levels. *MTARC1* mRNA levels were not significantly different between the two genotypes **(Fig. 2B)**. However, MTARC1 protein levels were significantly reduced in HLOs with the *MTARC1*-A165T (A/A) genotype compared to HLOs expressing *MTARC1*-WT (G/G) as revealed by Western blot (**Fig. 2C**) and confirmed with densitometric quantification of band intensities **(Fig. 2D)**.

Hepatic lipid accumulation is a key hallmark of MASLD development and progression (6). Previously, we defined a free fatty acid-induced steatosis/steatohepatitis HLO model by challenging HLOs with palmitic acid (PA) (31). We treated *MTARC1*-WT and *MTARC1*-A165T HLOs with PA and quantified steatosis using Bodipy staining (**Fig. 3A**). Following PA challenge, HLOs carrying the *MTARC1*-A165T variant exhibited a reduced Bodipy signal (Right panel) in comparison to *MTARC1*-WT HLOs (left panel), with consistent lipid phenotypes across all clones with the same genotype (**Fig. 3A**). Quantification of Bodipy intensities confirmed these observations (**Fig. 3B**). Under control conditions, a slight but non-significant decrease in Bodipy staining of *MATRC1*-A156T HLOs was also observed. To confirm the reduction of steatosis in HLOs carrying the protective variant, we next quantified intracellular triglyceride levels (**Fig. 3C**), which were also reduced in HLOs expressing the *MTARC1*-A165T variant under PA treatment. These findings show that *MTARC1*-A165T variant HLOs are protected from development of steatosis.

**Fig. 3.**
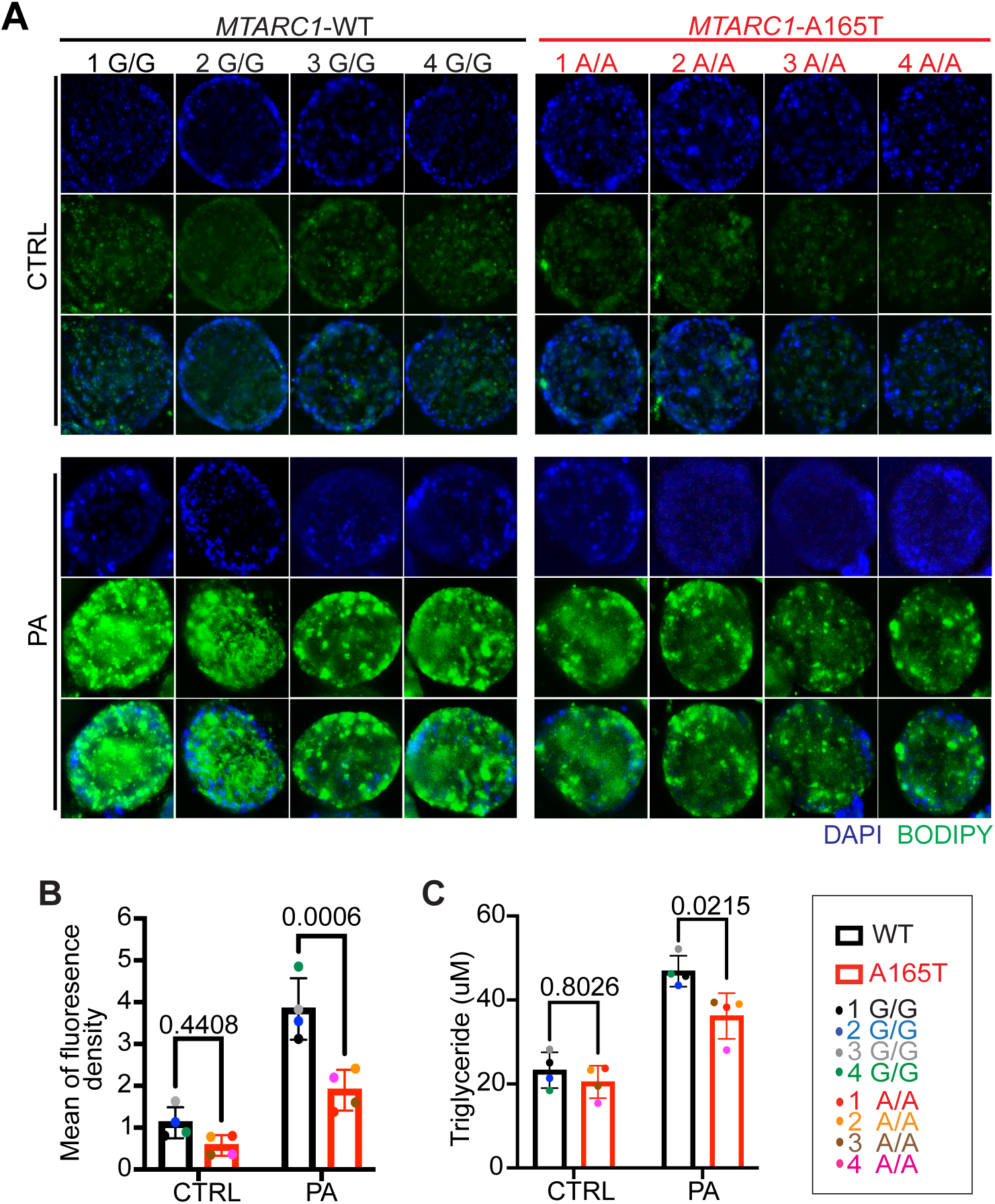
*MTARC1* A165T variant reduces lipid accumulation. (A) HLOs carrying *MTARC1*-WT and -A165T were treated with control condition or PA (500 uM) for 4 days on an orbital shaker, then stained with Bodipy (green) and Hoechst (blue). Images are representative of 3 independent experiments and in each experiment, every clone was differentiated in triplicate. Scale bar, 100 um. (B) The graph shows the mean fluorescence density which represents the mean fluorescence intensity of lipid droplets (Bodipy) normalized by organoid size. 15-19 HLOs per group. (C) Quantification of intracellular triglycerides in HLOs expressing *MTRAC1*-WT and -A165T. Presented data are from 3 independent experiments and in each experiment, every clone was differentiated in triplicate. Two-way ANOVA test was performed for statistical comparison for graphs B and C.

### *MTARC1* A165T variant reduces expression of genes in the de novo lipogenesis pathway

To further investigate how the *MTARC1*-A165T variant attenuated lipid accumulation in hepatocytes, we quantified the impact of the *MTARC1*-A165T variant on expression of genes that regulate lipid metabolism. *ACC1* and *FASN* are two key genes whose products regulate de novo lipogenesis by catalyzing the first two steps of fatty acid synthesis, respectively (37). Expression of *ACC1* was reduced in the *MTARC1*-A165T variant compared to *MTARC1*-WT HLOs in both standard culture conditions and with the addition of PA **(Fig. 4A).** *FASN* expression significantly decreased under standard conditions, but the addition of PA suppressed *FASN* to similar levels in both *MTARC1*-A165T variant and *MTARC1*-WT HLOs (**Fig. 4B**). We next quantified the expression of genes that regulate ß-oxidation: *CPT1a*, *CPT2* and *ACADAM*. *CPT1a* showed a trend towards reduced expression in the *MTARC1*-A165T variant compared to *MTARC1*-WT HLOs (**Fig. 4C)**, while *CPT2* and A*CADAM* expressions did not change **(Fig. 4D and 3E)**. Overall, these results suggest that the presence of the *MTARC1*-A165T variant may suppress steatosis in the HLO model by reducing de novo lipogenesis rather than increasing ß-oxidation.

**Fig. 4.**
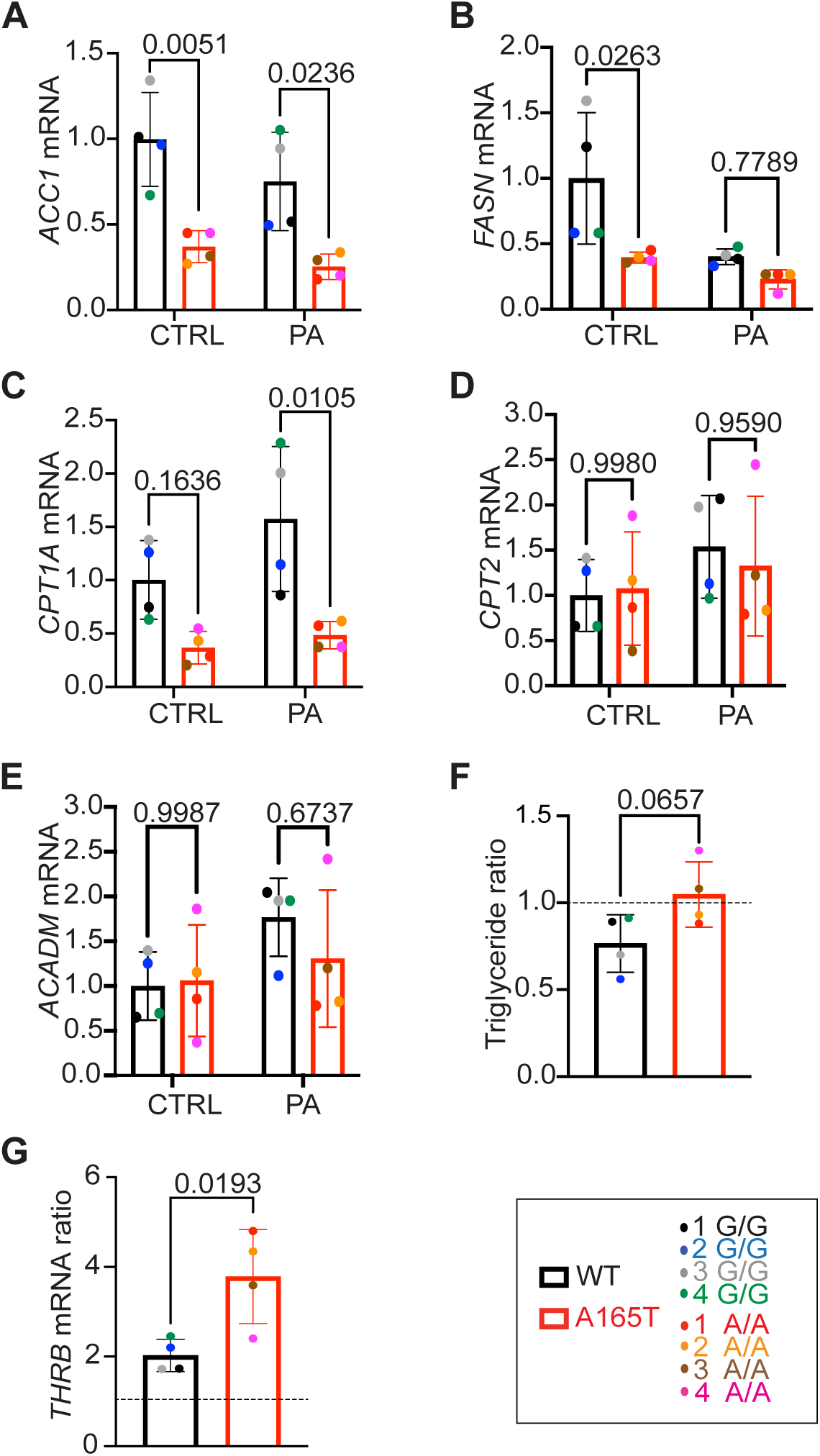
*MTARC1* A165T variant reduces expression of genes in the de novo lipogenesis pathway and attenuates the impact of resmetirom. *ACC1* (A), *FASN* (B), *CPT1a* (C)*, CPT2* (D)*, and ACADAM* (E) mRNA levels were quantified by RT-qPCR with *ACTB* serving as an endogenous control. Data are representative of 2 independent experiments, and in each experiment, every clone was differentiated in triplicate. Two-way ANOVA test was performed for statistical comparison for graphs A, B, C, D, and E. (F) *MTARC1*-WT and -A165T HLOs were treated with PA (500 uM) or the combination of PA (500 uM) and resmetirom (10 uM) for 4 days, then intracellular triglyceride was quantified. The graph represents intracellular triglyceride ratio quantified as the ratio of triglycerides in PA plus resmetirom compared to PA alone for each clone. Ratio < 1 reflects the reduction of triglycerides content in HLOs treated with resmetirom plus PA compared to PA alone. The dotted line indicates a ratio of 1. Data are included from 2 independent experiments, and in each experiment, every clone was differentiated in triplicate. P values were calculated using an unpaired two-tailed Student’s t-test. (G) *THRB* mRNA ratio was quantified in *MTARC1-*WT and A165T HLOs treated with PA or PA plus resmetirom. *ACTB* serves as an endogenous control. The graph represents the ratio of *THRB* mRNA level between PA plus resmetirom and PA alone. Ratio > 1, indicates the increase of *THRB* mRNA expression in PA plus resmetirom compared to PA alone. A dotted line indicates a ratio of 1. Data are included from 2 independent experiments, and in each experiment, every clone was differentiated in triplicate. P values were calculated using an unpaired two-tailed Student’s t-test.

### *MTARC1* A165T variant attenuates the impact of resmetirom

The THRB agonist, resmetirom, was recently approved for patients with MASH and can reduce steatosis in the setting of MASLD (8), and we wanted to understand how the *MTARC1*-A165T variant affects responses to resmetirom. We found that treatment of *MTARC1*-WT HLOs with resmetirom in the presence of PA suppressed triglyceride levels, while resmetirom did not further suppress triglyceride levels in HLOs expressing the *MTARC1*-A165T variant in the presence of PA **(Fig. 4F)**. While the *MTARC1*-A165T did not show a change in triglyceride levels in response to resmetirom, *THRB* expression was induced in both the variant and WT HLOs, consistent with resmetirom activity in both conditions **(Fig. 4G)**. These results show that while both variant and WT HLOs respond to resmetirom, resmetirom does not appear to provide any further benefit in lowering triglyceride levels in HLOs with the *MTARC1*-A165T variant.

### *MTARC1-* A165T decreases inflammation in HLOs

Inflammation is a key feature in MASLD progression from steatosis to steatohepatitis (38). To evaluate the impact of the *MTARC1*-A165T variant on the transition from steatosis to steatohepatitis, we quantified cytokine and chemokine expression in response to PA. We found that *MTARC1*-A165T variant HLOs secreted less IL-6, IL-1α, TNF-α and RANTES compared to WT HLOs. **(Fig. 5A-D)**. These results suggest that the A165T variant produces a muted inflammatory response, which could reduce the rate of progression from MASLD to MASH.

**Fig. 5.**
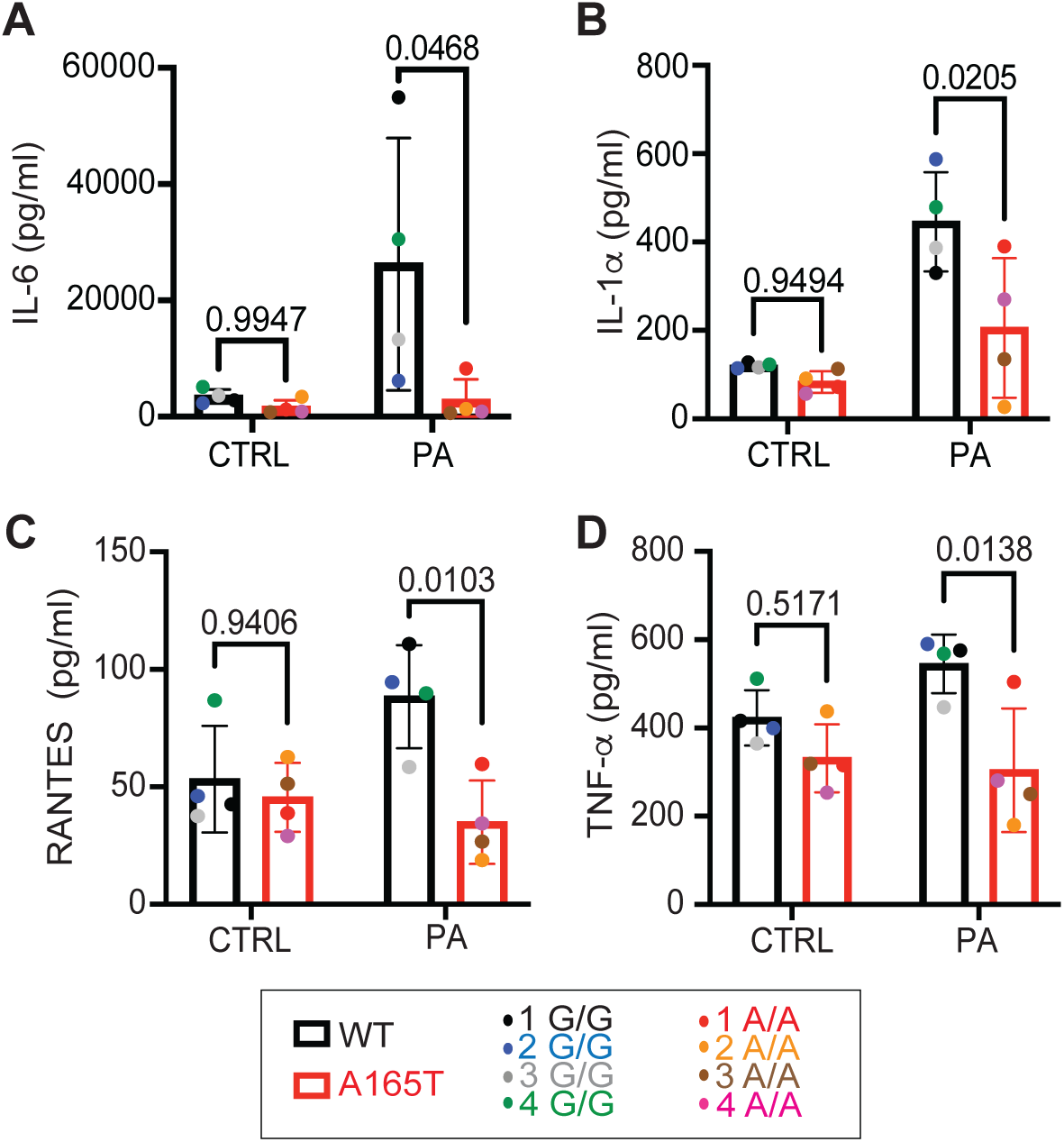
*MTARC1-* A165T decreases inflammation in HLOs. After PA treatment, media from *MTARC1*-WT and -A165T HLOs was analyzed to detect: IL-6 (A), IL-1α (B), TNF-α (C) and RANTES (D). Data are included from one experiment, and every clone was differentiated in triplicate. Two-way ANOVA test was performed for statistical comparison for all graphs.

### *MTARC1* A165T protects from liver fibrosis

In MASLD, hepatic stellate cells (HSCs) play a central role in the development of fibrosis and disease progression. Once activated, HSCs produce the extracellular matrix (ECM) proteins that accumulate in fibrosis (39). The presence of HSCs in our HLOs culture enables liver fibrosis modeling using TGF-β1 treatment (31). We evaluated the fibrotic response of HLOs carrying *MTARC1*-WT and *MTARC1*-A165T variants to treatment with TGF-β1. *ACTA2* expression, which marks HSC activation (39), was reduced in HLOs expressing the *MTARC1*-A165T variant (**Fig 6A-E**). Additionally, a significant reduction in *COL1A1* and *COL1A2* mRNA levels was observed in HLOs carrying *MTARC1*-165T variant compared to *MTARC1*-WT HLOs (**Fig. 6F -G**), and type 1 collagen protein levels were also reduced in *MTARC1*-165T HLOs (**Fig. 6H-K**). These results show that the *MTARC1*-165T variant suppresses HSC activation and is associated with reduced expression of type I collagen, a key component of the ECM contributing to the fibrotic scar.

**Fig. 6.**
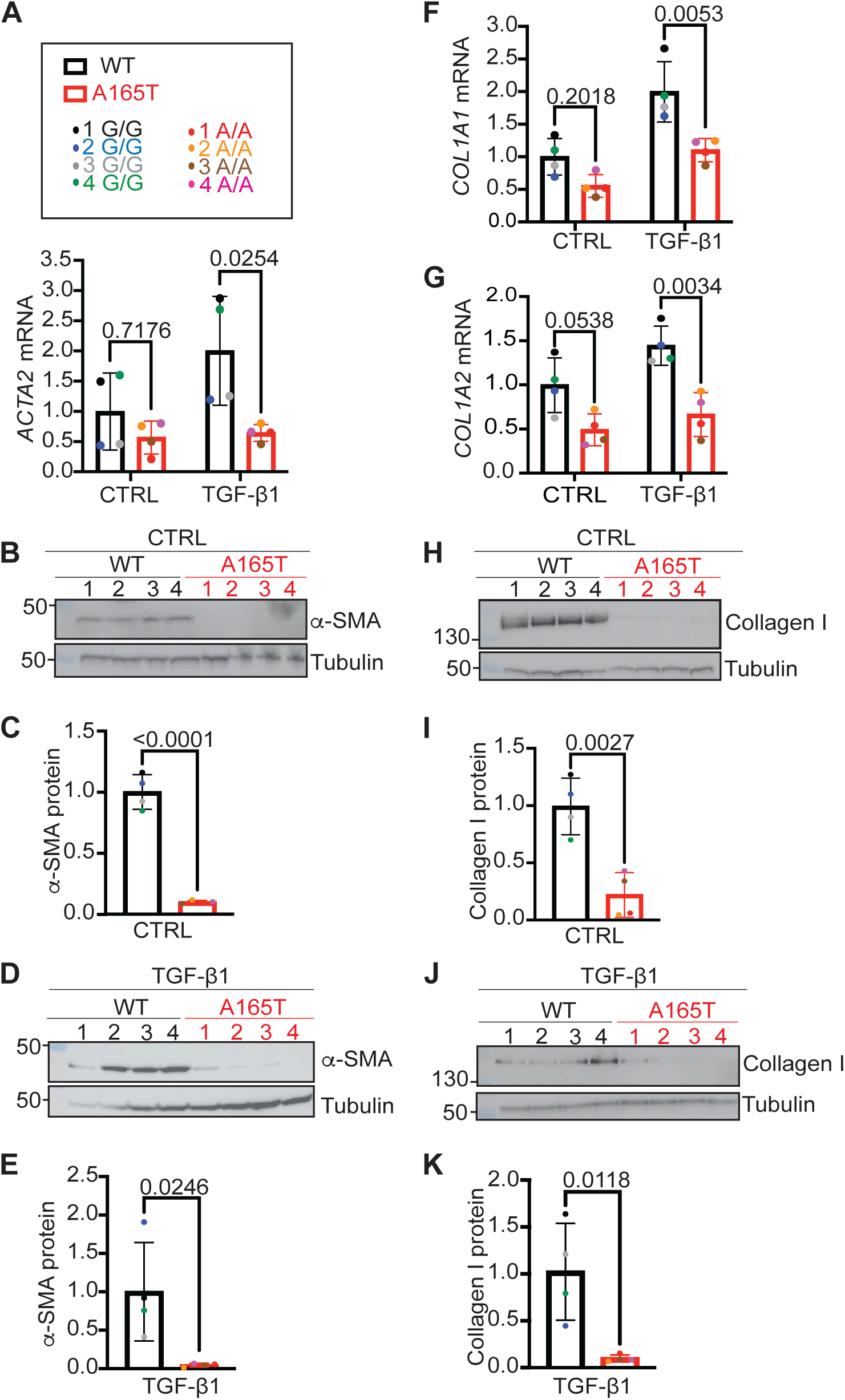
*MTARC1* A165T variant protects from liver fibrosis. *MTARC1*-WT and -A165T HLOs were treated with control condition or TGF-β1 (10 ng/mL) for 4 days on an orbital shaker before analysis. (A) *ACTA2* was quantified by RT-qPCR with *ACTB* serving as an endogenous control. (B) Western blot of α-SMA protein (encoded by *ACTA2*) levels in HLOs under control conditions. (C) Quantification of α-SMA signal shown in B. (D) Western blot of α-SMA protein levels after TGF-β1, and (E) represents the quantification of D. (F) *COL1A1* levels were quantified by RT-qPCR with *ACTB* serving as an endogenous control. (G) Quantification of *COL1A2* mRNA levels with *ACTB* serving as an endogenous control. (H) Collagen type I protein levels detected by Western blot under control culture conditions. (I) Quantification of Collagen type I protein from (H). (J) Collagen type I protein levels detected by Western blot after TGF-β1 treatment. (K) Quantification of Collagen type I protein from (J). Two-way ANOVA tests were performed for statistical analyses of data in B, G and H and unpaired two-tailed Student’s test was used for data in D, F, J and L. Data are representative of 3 independent experiments, and in each experiment, every clone was differentiated in triplicate.

## DISCUSSION

The human *MTARC1* gene was discovered in 2006 and has been linked to drug metabolism and detoxification (24). With the discovery of the protective *MTARC1 rs2642438* (p.A165T) variant (18), there has been growing interest directed toward understanding the role of *MTARC1* and how protective variants modulate the development and progression of chronic liver disease.

Current models have yielded limited insights into the functions of *MTARC1* and its variants. In vivo studies have reported contradictory results regarding *Mtarc1* function, with some studies showing that loss of *Mtarc1* did not affect liver steatosis or fibrosis and others showing a protective effect in MASH models (30, 40). Furthermore, immortalized and primary hepatocyte models present limitations and fail to replicate the complex functions of the liver and have relied on ectopic expression or depletion of *MTARC1* (*26, 41*). In recent years, organoids derived from pluripotent cells have emerged as a promising and powerful alternative to study human liver diseases in culture (42). With their 3D multicellular organization, HLOs can better approximate the cellular heterogeneity, cell crosstalk, and functions of the liver. In the current study, we generated an HLO model to study the *MTARC1*-A165T variant by performing prime editing in hPSCs to create *MTARC1*-A165T homozygous and *MTARC1*-WT homozygous hPSCs from hPSCs originally expressing one variant and one WT allele. By analyzing HLOs created from these hPSCs, we can study the impact of a single variant on an otherwise shared genetic background.

The *MTARC1*-165T variant is linked to decreased protein levels, as observed in hepatocarcinoma cell lines and mouse livers overexpressing the human *MTARC1* A165T variant (26, 30, 40, 43). These findings have been supported by studies showing that the MTARC1 A165T protein exhibits reduced stability, greater ubiquitination, and accelerated degradation by proteasome compared to MTARC1 WT protein (27, 44). Data to address protein stability from human samples have been more limited, with studies from deceased donors showing reduced MTARC1 protein levels in the presence of the *MTARC1* A165T variant (40) but studies from pediatric liver biopsies showing no significant difference (45). Our findings in HLOs further support the conclusion that the *MTARC1*-A165T is associated with reduced MTARC1 protein levels despite similar levels of *MTARC1* transcripts.

The *MTARC1*-165T variant is also linked to reduced steatosis in patients (18, 45), and these observations have been extended to human primary hepatocytes and mouse models through depletion of *MTARC1* (28–30). Consistent with these findings, we observed that HLOs carrying *MTARC1*-A165T variant were protected from lipid/triglyceride accumulation when challenged with PA. We also found that the reduction of lipids in *MTARC1*-A165T variant HLOs is associated with lower expression of genes that regulate the de novo lipogenesis *(ACC1* and *FASN*). While increased ß-oxidation has been linked to the A165T variant in primary hepatocytes (41)*, CPT2* and *ACADAM* levels did not change significantly in the A165T variant HLOs, and *CPT1* even showed a decreasing trend in the variant, all pointing away from a role for increased ß-oxidation as a path to reduce lipid accumulation in the HLO model. These findings suggest a central role of *MTARC1* in lipid handling, but further mechanistic studies are needed to clarify its function.

Consistent with clinical observations (23), we found that HLOs expressing the *MTARC1* A165T variant produce lower levels of cytokines and chemokines in response to PA treatment. *MTARC1* expression is restricted to hepatocytes (25), and our HLO model does not contain macrophages (31), suggesting that inflammatory signals produced by hepatocytes to recruit macrophages (46) are also suppressed in the A165T variant. Additionally, IL-6, IL-1α and TNF-α can promote HSC activation and collagen production (47) and decreased expression of these ligands may also be the path by which the A165T variant reduces fibrogenesis, even in the presence of TGF-ß1.

Resmetirom provides a significant advance in treatment options for patients with MASH (10, 39), however there are not yet studies that evaluate how SNPs linked to disease progression may impact responses to resmetirom therapy. Our data show that the decrease in triglyceride levels observed in the *MTARC1*-A165T variant HLOs is not further reduced with resmetirom treatment, raising the possibility that patients with the *MTARC*1-A165T variant may benefit less from this treatment. Resmetirom was also observed to be less effective in reducing steatosis in human hepatocytes containing the I148M (disease) variant of PNPLA3 compared to human hepatocytes containing the reference sequence, in a microphysiology system (48). These findings suggest that understanding the impact of disease-related SNPs may help select patients who are more likely to benefit from resmetirom or future drugs to treat MASH.

Establishing HLOs from hPSCs engineered to contain disease-relevant SNPs provides a robust platform for investigating genetic mechanisms that shape liver disease. In this study, we modeled the *MTARC1*-A165T variant and found it to be associated with reduced MTARC1 protein expression and protection against lipid accumulation, linked to suppression of de novo lipogenesis. We also observed decreased secretion of inflammatory cytokines and chemokines, which may contribute to the attenuated fibrotic response seen following TGF-β1 stimulation. Notably, the variant lowers triglyceride levels to such an extent that resmetirom treatment did not confer additional benefit, suggesting that genetic background may influence therapeutic efficacy. Collectively, these findings underscore the utility of engineered HLOs for dissecting disease-relevant pathways and highlight the importance of *MTARC1* in advancing our understanding and treatment of chronic liver disease.

## METHODS

### hPSCs culture, differentiation into HLOs and liver injury induction

H1 human pluripotent stem cells (WiCell, WA01- NIH registration number 0043) were cultured and differentiated into HLOs as previously described (31, 32). In brief, hPSCs were maintained in mTeSR plus medium (StemCell Technologies, 100-0276) in plates pre-coated with Matrigel (Corning, 354230). After reaching confluency, hPSCs were differentiated to definitive endoderm then to foregut spheroids. On day 7 of differentiation, cells were embedded in Matrigel and treated for 4 days with retinoic acid (Tocris, 60695) before switching to hepatocyte culture medium (Lonza, CC-3198) (33). On day 21, HLOs were extracted from Matrigel and cultured on an orbital shaker. To induce liver injury, isolated HLOs were treated for 4 days with palmitic acid (PA, 500 uM) or transforming growth factor-beta 1 (TGF-β1, 10 ng/mL).

### Prime editing

To generate the *MTARC1* -wildtype (WT) and -A165T clones, hPSCs were edited using prime editing strategy 5 (PE5) as described in (34, 35). Briefly, hPSCs were detached using accutase (StemCell Technologies, 07920) and nucleofected with 1 ug of in vitro transcribed Cas9-nickase fused to reverse transcriptase (*nCAS9-RT*) mRNA, 1 ug of in vitro transcribed engineered dominant-negative MLH1 mutant (*MLH1dn*) mRNA, 100 pmol of chemically modified synthetic engineered pegRNA (epegRNA) and 60 pmol of synthetic nicking single guide RNA (sgRNA) using the P3 Primary cell 4D Nucleofector X kit S (Lonza, V4XP-3032). 72 hours after the first nucleofection, hPSCs were nucleofected a second time using the same reagents. Edited hPSCs were cultured for additional 72 hours to recover and then single clones were picked, cultured in 96 well plates until confluency and screened for the targeted edit. Genomic DNA from single-cell-derived colonies was isolated using the Arcturus PicoPure DNA Extraction kit (Applied biosystems, 11815-00), PCR amplified using primers designed to amplify around the target edit (primer sequences are attached in Table 1) and amplicons were subjected to Sanger sequencing. In vitro transcribed *nCAS9-RT* and *MLH1dn* mRNAs were prepared as described in (34). epegRNAs and sgRNA (sequences are included in Table S1) were designed using the pegFinder website (36).

### Quantitative reverse transcriptase real-time PCR (RT-qPCR)

RNA was isolated using TRIzol reagent (Invitrogen, 15596026), reverse transcribed using the iScript gDNA clear cDNA synthesis kit (Bio-Rad, 1725035) according to manufacturer’s instructions. RT-qPCR was performed using SYBR Green universal master mix (Applied Biosystems, 4309155). Actin Beta (*ACTB)* was used as an internal control for all the experiments. All used primer sequences are listed in Table S2.

### Pluripotent marker staining

hPSCs were fixed with 4% paraformaldehyde (Thermo Scientific, J61899-AK) for 10 min, permeabilized using 0.1% Triton, blocked for 1 hour in 1% Bovine Serum Albumin (BSA) solution (Sigma, A4737) and then stained with antibodies recognizing OCT4 and NANOG (Table S3) overnight at 4 °C. The following day, cells were incubated with Donkey anti-Goat IgG Cross-Adsorbed Secondary antibodies Alexa Fluor 488 and 594 and DAPI and then were visualized and imaged with the Leica DMi8 automated Microscope (Leica Systems) using the 10x objective.

### Western blots

HLOs were lysed in RIPA Buffer (Boston BioProducts, BP-115D) supplemented with Halt protease inhibitor cocktail (Thermo Scientific, 78438). After homogenization, the lysates were centrifuged at 20,000 g at 4 °C for 10 min. Supernatants were then collected. Protein concentrations were quantified using Pierce BCA protein assay kit (Thermo Scientific, 23227), and equal protein quantities supplemented with Bolt LDS Sample Buffer (Invitrogen, cat# B0007) and Bolt Sample Reducing Agent (Invitrogen, cat# B0009) were loaded in Bolt 4-12% Bis-Tris gels (Invitrogen, NW04120BOX). After protein transfer using iBlot 2 Dry Blotting system, membranes were blocked with 3% milk (Lab Scientific, M0841) in Tris Buffered Saline Tween (TBST) (Boston BioProducts, IBB-180X) for 1 hour at room temperature and incubated overnight with primary antibodies at 4 °C. The following day, membranes were washed with TBST, incubated with secondary antibodies for 1 hour, washed again, incubated with the Super Signal West Pico PLUS chemiluminescent substrates (Thermo Scientific, 34580) and scanned using a LI-COR Odyssey M imaging machine. All antibodies used are listed in Table S3. Tubulin was used as a loading control. Densitometry was performed using Image J- Fiji software.

### Bodipy staining

HLOs cultured in control l or PA conditions were collected and washed with PBS. Lipid droplets were stained with Bodipy 493/503 and nuclei with DAPI for 40 min at 37°C. The stained HLOs were then visualized and imaged with the Leica DMi8 automated Microscope (Leica Systems) using the 10× objective. Image analyses and fluorescence intensity quantification were performed using the ImageJ-Fiji software.

### Triglyceride assay

HLOs were homogenized in 4% NP-40 solution (Fisher Scientific, PI85124). Triglyceride levels were then measured using a fluorometric triglyceride quantification assay kit (Abcam, Ab65336) according to the manufacturer’s instructions.

### Cytokines and chemokines quantification

HLOs were cultured in control or PA conditions. Media were collected, and secreted cytokines and chemokines were quantified using the customized milliplex human cytokine/chemokine multiplex assay (Millipore, HCYTA-60K) following the manufacturer’s recommendations.

### Statistics

All graphs and statistical analyses were performed using GraphPad Prism 10. Unpaired two-tailed Student’s test and two-way ANOVA test were used for statistical analyses as indicated. Error bars indicate standard deviation (SD).

## AUTHOR CONTRIBUTIONS

A.C.M and A.B.S conceived and designed the study. A.B.S performed the experiments and data analysis with assistance from A.W, S.D.G, N.A, B.T, S.T, S.S, and A.C. The manuscript was written by A.B.S and A.C.M with input from all other authors.

## ACKNOWLEDGEMENTS

We would like to thank Erik Sontheimer and Zexiang Chen for sharing their expertise and protocols for prime editing. This work was supported by internal funding from UMass Chan Medical School.

## ABBREVIATIONS

Chronic liver disease (CLD), Metabolic dysfunction-associated steatotic liver disease (MASLD), Metabolic dysfunction-associated steatohepatitis (MASH), Genome-wide association studies (GWAS), Single nucleotide polymorphism (SNP), Mitochondrial amidoxime reducing component 1 (*MTARC1),* Human pluripotent stem cells (hPSCs), Human liver organoids (HLOs), Thyroid hormone receptor-beta (THRB), Patatin-like phospholipase domain-containing 3 (*PNPLA3*), Transmembrane 6 superfamily member 2 (*TM6SF2*), Glucokinase regulatory protein (*GCKR*), Membrane-bound O-acyltransferase domain containing 7 (*MBOAT7*), Hydroxysteroid 17 beta dehydrogenase 13 (*HSD17B13*), Alanine transaminase (ALT), Aspartate transaminase (AST), N-acetylgalactosamine conjugated siRNAs (GalNac-siRNA), Wildtype (WT), Cas9-nickase fused to reverse transcriptase (nCAS9-RT), Engineered pegRNA (epegRNA), Single guide RNA (sgRNA), Actin Beta (*ACTB),* Palmitic acid (PA), Transforming growth factor-beta 1 (TGF-β1), Hepatic stellate cells (HSCs), Interleukin-6 (IL-6), Interleukin-1alpha (IL-1α), Tumor necrosis factor-alpha (TNF-α), Regulated upon activation normal T cell expressed and secreted (RANTES), Alpha-smooth muscle actin (α-SMA), Standard deviation (SD).

